# Leishmaniinae: evolutionary inferences based on protein expression profiles (PhyloQuant) congruent with phylogenetic relationships among *Leishmania, Endotrypanum, Porcisia, Zelonia, Crithidia,* and *Leptomonas*

**DOI:** 10.1101/2023.09.22.558958

**Authors:** Simon Ngao Mule, Evaristo Villalba Alemán, Livia Rosa Fernandes, Joyce S. Saad, Gilberto Santos de Oliveira, Deivid Martins, Claudia Blanes Angeli, Deborah Brandt-Almeida, Mauro Cortez, Martin Røssel Larsen, Jeffrey J. Shaw, Marta M. G. Teixeira, Giuseppe Palmisano

## Abstract

Evolutionary relationships among parasites of the subfamily Leishmaniinae, which comprises pathogen agents of leishmaniasis, were inferred based on differential protein expression profiles from mass spectrometry-based quantitative data using the PhyloQuant method. Evolutionary distances following identification and quantification of protein and peptide abundances using Proteome Discoverer (PD) and MaxQuant (MQ) softwares were estimated for 11 species from 6 Leishmaniinae genera. Results clustered all dixenous species of the genus *Leishmania*, subgenera *L. (Leishmania), L. (Viannia)* and *L. (Mundinia),* sister to the dixenous species of genera *Endotrypanum* and *Porcisia.* Positioned basal to the assemblage formed by all these parasites were the species of genera *Zelonia*, *Crithidia* and *Leptomonas*, so far described as monoxenous of insects although eventually reported from humans. Inferences based on protein expression profiles were congruent with currently established phylogeny using DNA sequences. Our results reinforce PhyloQuant as a valuable approach to infer evolutionary relationships consistent with genera, subgenera, and species-specific biological characteristics, able to resolve within Leishmaniinae, which is comprised of very tightly related trypanosomatids that are just beginning to be phylogenetically unravelled. In additional to evolutionary history, mapping of species-specific protein expression is paramount to understand differences in infection processes, disease presentations, tissue tropisms, potential to jump from insects to vertebrates including humans, and potential targets for species-specific diagnostic and drug development.

## Introduction

The PhyloQuant approach was developed to infer evolutionary relationships between organisms based on systems-wide differential protein expression obtained from mass spectrometry data [1]. This method was first applied to *Trypanosoma cruzi* and close-related bat trypanosomes of the subgenus *Schizotrypanum*, all species nesting into the major clade *T. cruzi* [2]. We demonstrated that PhyloQuant can arrange these trypanosomes according to species and intraspecific classification of *T. cruzi* DTUs (Discrete Typing Units TcI-TcVII), consistent with phylogeny based on multiple DNA sequences. Moreover, strains, DTUs, and species-specific proteins and protein expression profiles were highlighted by PhyloQuant, providing relevant data to any attempts in elucidating aspects on host-parasite interaction, adaptation to new hosts and vectors, and the evolution of pathogenicity within the clade *T. cruzi* [1].

Trypanosomatids are obligatory parasites usually classified as monoxenous, those that develop in a single host mostly insects, and dixenous species developing in vertebrates or plants with insects serving as vectors. In the current study, we applied the PhyloQuant method to infer evolutionary relationships of trypanosomatids nested into the subfamily Leishmaniinae. This subfamily comprises both dixenous species of *Leishmania*, *Endotrypanum* and *Porcisia*, majority being transmitted by sand flies, and monoxenous species of genera *Zelonia, Novymonas* and *Borovskyia* (parasites of dipterans and heteropterans), and genera *Leptomonas, Lotmaria* and *Crithidia* (dipterans, heteropterans and hymenopterans) forming the most basal group of the subfamily [3–5].

*Leishmania* species are responsible for a broad spectrum of neglected tropical diseases collectively termed leishmaniasis, affecting over 12 million people in more than 98 countries where more than one billion people are at risk of contracting the diseases. Depending on the species of *Leishmania*, host genetics and immune status, the infection evolved to clinical forms referred as to cutaneous (localized, diffused, or disseminated) and visceral leishmaniasis. Visceral leishmaniasis causes > 20000 deaths annually while cutaneous leishmaniasis affects the life quality of millions [6]. The *genus Leishmania* is composed of four subgenera: *L.* (*Leishmania*), *L*. (*Viannia*), *L.* (*Sauroleishmania*) and *L.* (*Mundinia*) [4, 5, 7]. Of medical importance are more than 20 species of *L. (Leishmania)* and *L. (Viannia*), which are agents of visceral, cutaneous, and mucocutaneous disease. However, an increasing number of species of *L.* (*Mundinia*), likely transmitted by biting midges (Ceratopogonidae) have been reported in diverse mammal hosts including humans. Both *L.* (*Leishmania*) and *L*. (*Mundinia*) are distributed throughout the Old and New Worlds, whereas *Leishmania* (*Viannia)*, *Endotrypanum*, and *Porcisia* are exclusive to the New World [8–12].

Phylogenetic studies have demonstrated that in addition to *Leishmania* spp., human infection can be caused by *Endotrypanum colombiensis,* a former species of *Leishmania* transferred to *Endotrypanum*, whose species are in general transmitted by sandflies to sloths, the main hosts of this genus [4, 13–16]. The species of *Porcisia*, so far exclusive to Neotropical porcupines, form together with the genus *Endotrypanum* a clade sister to *Leishmania*. The life cycle of *Porcisia* remains unclear, but a recent experimental study demonstrated an unexpected contaminative transmission of *P. deanei* by both sand flies and biting midges [17]. Zelonia are apparently a monoxenous genus consistently placed in the major clade nesting all dixenous species. Species of Zelonia were reported in Central to South America, from hemipterans and dipterans, while *Zelonia australiensis,* from black fly, was reported in Australia [4, 18]. With the availability of molecular approaches to identify trypanosomatids in general, including mixed infections, *Leishmania* have been associated to concomitant infection with *L. seymouri,* as discovered recently in humans from India and Sri Lanka [19–22], and *Crithidia* sp. identified in fatal cases of visceral disease and, recently, in a case of concomitant visceral and cutaneous leishmaniasis in Brazil [23, 24].

Traditional approaches for the identification of trypanosomatids causing leishmaniasis based on clinical disease, serology, zymodemes and *Leishmania-*specific conventional PCRs are insufficient to assess the increasing genetic diversity within the subfamily Leishmaniinae. Currently, the identification of trypanosomatids requires phylogenetic inferences including pathogenic and closely related but apparently non-pathogenic species. Phylogenetic and taxonomic controversies together with underestimated genetic diversity, enigmatic nature of some species, and human cases of usually non-infective species to humans prompted us to use a proteomic approach to contribute to a broader understanding of Leishmaniinae. Using the PhyloQuant approach, here we benchmarked the use of quantitative proteomics and differential protein expression to infer evolutionary relationships between 11 species from six genera of Leishmaniinae. The data were consistent with accepted phylogeny based on gene sequences, provide the first proteomes of *Endotrypanum monterogeii*, *Crithidia thermophila*, *Zelonia costaricensis*, and *Leptomonas moramango*, and highlight species-specific regulated proteins with possible correlation to species-specific biological traits of host-parasite-vector interactions, human infectivity, and pathogenicity.

## Material and Methods

### Parasite cultures

Representative species in the subfamily Leishmaniinae analyzed in this study included *L. (L.) amazonensis*, *L. (L.) infantum chagasi, L. (V.) braziliensis* and *L. (M.) enrietti, P. deanei*, *E. schaudinni (herein called E. monterogeii)*, *Z. costaricensis*, *Zelonia sp.*, *C. thermophila*, *L. seymouri*, and *L. moramango.* Their respective hosts and localities of origin are summarized in **Table 1**. *L. (Leishmania)* and *L. (Viannia)* species were cultivated in M199 medium supplemented with 20% fetal bovine serum (FBS) and 2% sterilized human urine, while the other species were cultured in TC 100 medium supplemented with 10% FBS. All the species were cultured at 28°C. The parasites were cultured in biological triplicates and collected at the exponential growth phase of promastigote forms by centrifugation at 2100 g for 10 min. The pellets were washed three times with 1× PBS (137mM NaCl, 2.7 mM KCl, 10 mM Na_2_HPO_4_, 1.8 mM of KH_2_PO_4_, pH 7.4) before subsequent analysis. The parasites included in the present study are deposited in the TCC-USP (Trypanosomatid Culture Collection, Department of Parasitology, University of São Paulo).

**Table 1.**
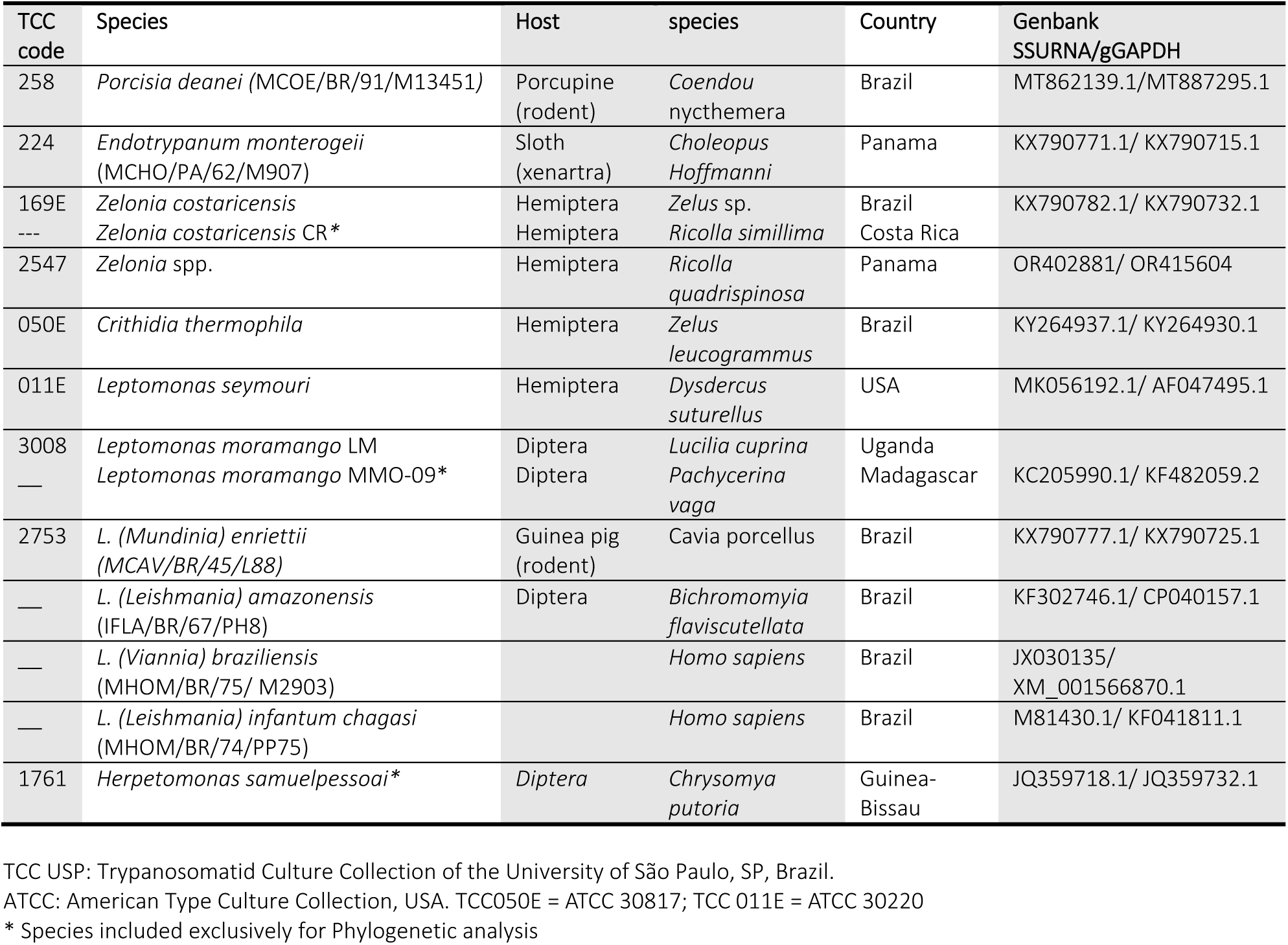
Trypanosomatids included in PhyloQuant and/or phylogenetic analyses.

### Protein extraction and sample preparation

Parasite cells were lysed in 8M urea supplemented with a 1X Protease inhibitor cocktail (cOmplete™, Mini, EDTA-free Protease Inhibitor Cocktail). The cells were lysed by five freeze-thaw cycles, and the proteins were quantified using a Qubit fluorometer. Protein disulphide bonds were reduced using 10 mM DTT by incubation for 45 min at 30ᵒC before alkylation at room temperature using 40 mM iodoacetamide for 30 min in the dark. The alkylation reaction was quenched using 5 mM DTT, and the lysis buffer was diluted ten times with 50 mM NH_4_CO_3_. The proteins were subsequently digested overnight with molecular grade trypsin enzyme (Promega) using a ratio of 1:50µg (w/w) enzyme to proteins at 30°C, acidified with 1% (v/v) trifluoroacetic acid, desalted by reverse phase (RP) using C18 in-house columns and dried down by vacuum centrifugation. Resultant tryptic peptides were submitted to nano LC-MSMS analysis using the following parameters described below.

### nLC-MS/MS analysis

The proteomes of the trypanosome species included in this study were analysed by liquid chromatography coupled with tandem mass spectrometry. *L.* (*L.*) *amazonensis*, *L.* (*V.*) *braziliensis* and *L.* (*L.*) *infantum chagasi* were analyzed separately, but using the same nLC-MS/MS equipment as the other species in this study. Liquid chromatography was performed on a Reprosil®-Pur C18-aQ nano HPLC column (3µm; Dr. Maisch GmbH, Germany) using Easy-LC nano-HPLC (Proxeon, Odense, Denmark) with a nanospray source (Thermo Scientific) mass spectrometer Q Exactive™ HF hybrid quadrupole-Orbitrap (Thermo Scientific). The HPLC gradient was 0–34% solvent B (A 0.1% formic acid; B 90% ACN, 0.1% formic acid) at a flowrate of 300 nL/min. The precursor ions were generated in positive ion-mode and selected in data-dependent acquisition. Run time of 130 (*L. (Leishmania)* and *L. (Viannia)*) or 100 min (the other 8 species), with a 400-1600 (*L. (Leishmania)* and *L. (Viannia)*) or 350-1600 (others) m/z scan range at a resolution of 120,000 FWHM, target AGC of 3×10^6^ ions, and maximum injection time of 100 ms were used. The MS/MS spectra of the 20 most abundant peptides were obtained by higher-energy collision dissociation (HCD) with acquisition in the orbitrap at a resolution of 30,000 FWHM, isolation window: 1.2 m/z, scan range of 200 – 2000, target AGC of 1×10^5^ ions; dynamic exclusion of 25 s (*L. (Leishmania)* and *L. (Viannia)*) and 30 s (others); intensity threshold was set to 2×10^4^. The mass spectrometry proteomics data have been deposited to the ProteomeXchange Consortium via the PRIDE [25] partner repository with the dataset identifier PXD044195.

### Database searches, protein, and peptide quantification

Identification and label-free (LFQ) quantification of proteins and peptides from the LC-MS/MS raw files was performed using two software for subsequent comparison; MaxQuant (MQ) (v 1.6.15.0) [26] and Proteome Discoverer (PD) (v 2.5), Thermo Scientific. For all analysis, MS/MS spectra were searched against the combined Uniprot *L. mexicana*, *L. braziliensis*, *L. infantum, L. seymouri* (ATCC 30220), *P. hertigi* (C119), and *E. monterogeii* database (48,766 entries; downloaded 31.03.2022) concatenated with a common contaminants protein database. For MaxQuant searches, the raw data were searched with a PSM and protein FDR less than 1%, MS/MS tolerance (FTMS) of 20 ppm and MS/MS tolerance (ITMS) of 0.5 Da. Two missed cleavages were allowed for trypsin cleavage. Carbamidomethylation of cysteine (57.0215 Da) was set as a fixed modification, while oxidation of methionine (15.9949 Da), deamidation NQ (+ 0.9840 Da) and protein N-terminal acetylation (42.0105 Da) were set as variable modifications. The match between runs (MBR) feature of MaxQuant was analysed both as activated or deactivated for identification comparisons. Match between runs was applied with a 0.7 min match time window and a 20 min alignment time window. Identified proteins were filtered to achieve a protein false discovery rate (FDR) of less than 1%. Intensity-based absolute quantification (iBAQ) and label-free quantification (LFQ) intensities were used for subsequent analyses, filtering for proteins identified with ≥2 peptides.

For Proteome Discoverer analysis, the Sequest HT processing node was used for database search. Trypsin digestion was set with two missed cleavages, ten (10) ppm precursor ion tolerance and 0.05 Da fragment ions mass tolerance. Carbamidomethylation of cysteine was set as a fixed modification, while methionine oxidation, asparagine, and glutamine deamidation and protein N-terminal acetylation were set as dynamic/variable modifications. Percolator, peptide validator and protein FDR validator nodes were used to filter PSMs, peptides, and proteins with FDR < 0.01. Label-free quantification was performed using the Minora algorithm of the processing workflow using minimal trace length of 5, max ΔRT of isotope pattern multiplets of 0.2 min, and high confidence PSMs. Precursor ions quantifier node and the Feature mapper were included in the consensus workflow to allow for retention time alignment (MBR). The highest scoring proteins (master proteins) and proteins identified with ≥2 peptides were exclusively used for subsequent analyses.

Proteins identified and quantified by both softwares were further processed using Perseus (v.1.6.15.0) [27] for principal component analysis, imputation of missing values, and statistical analysis. Contaminants, proteins only identified by site, proteins identified in the reverse database, and proteins identified with ≤ 1 peptides were removed before further analyses. Unsupervised Analysis of variance (ANOVA) was applied to identify the differentially expressed proteins and peptides between the protozoan species under study with Benjamini-Hochberg-based FDR correction at a FDR<0.05. Euclidean distance-based hierarchical clustering of the samples based on statistically significant differentially expressed proteins was performed following Z-score normalization.

### Evolutionary inference using the PhyloQuant method

The PhyloQuant method was used to infer the evolutionary relationships of the parasites under study as previously described [1]. Briefly, intensity-based absolute quantification (iBAQ) values were evaluated both as normalized and non-normalized quantitative features. Normalized intensities for label free protein and peptide quantification using MQ and PD searches were also included for evolutionary inference. TNT [28] (sponsored by Willi Hennig Society) was used for evolutionary inference. Bootstrap support values above 50% were illustrated on the evolutionary trees. TNT was also used to map the synapomorphies supporting each species or monophyletic assemblage. Expression profiles were visualized by a heatmap following statistical analysis and Euclidean distance clustering.

### Phylogenetic analysis and Mantel test correlation

For comparison with PhyloQuant evolutionary relationships, we phylogenetically analysed the organisms under study based on concatenated SSUrRNA and gGAPDH sequences as reported previously [3–5]. All sequences are available from GenBank, and accession numbers are showed on **Table 1**. Both SSU rRNA and GAPDH sequences of *L. moramango* LM from Uganda were identical to sequences previously described for *L. moramango* MMO-09 from Madagascar. Similarly, sequences from *L. enriettii* TCC2114 and *L. enriettii* TCC2753 were identical. *L. (Leishmania) amazonensis* isolate LaPH8.rRNA.01 18S ribosomal RNA and *L. (Leishmania) amazonensis* UA301 gGAPDH were identical to *L. (Leishmania) amazonensis* (IFLA/BR/67/PH8), SSU rRNA for *Leishmania braziliensis* isolate LbrET.rRNA.01 was identical to *L. (Viannia) braziliensis* (MHOM/BR/75/ M2903), and *Leishmania chagasi* isolate CBT 13 glycosomal glyceraldehyde 3-phosphate dehydrogenase (gGAPDH) gene was identical to *L. (Leishmania) chagasi* (MHOM/BR/1972/LD). SSU rRNA and gGAPDH genes for Zelonia spp. TCC2547 were amplified by PCR and sequenced, and have been deposited in GenBank under the accession numbers OR402881 and OR415604, respectively (**Table 1**). Evolutionary trees from PhyloQuant and sequence-based phylogenetic analysis were evaluated for levels of congruence using the Mantel test [29]. The ‘ape’ (analysis of phylogenetics and evolution) package [30] in the R statistical environment was used to calculate the levels of correlation using 1000 permutations. Ten random tree topologies sharing similar node names used in this study were included for comparison.

### Synapomorphies and analysis of species-specific differentially upregulated proteins

TNT was used to map the synapomorphies supporting each node/clade for one of the PhyloQuant trees generated earlier with high mantel test scores. Moreover, Euclidean distance-based clustering of differentially expressed proteins was performed in the Perseus software. Regulated proteins for each species under study were selected following ANOVA with Benjamini-Hochberg-based FDR correction at a FDR<0.05. Synapomorphies and species-specific upregulated proteins for the species under study were highlighted based on regulated proteins identified and quantified using PD software after 70 % valid value cut-off.

## Results

To benchmark the PhyloQuant method for evolutionary inference and taxonomic resolution, we analysed a set of highly closely related trypanosomatids nested in the *Leishmaniinae* subfamily using different software and mass spectrometry-based quantitative features (**Figure 1**). A total of 11 species with different host species, life cycle, geographical distribution, and clinical outcomes were compared (**Table 1**). In this study, we report for the first time the proteomes of *L. moramango*, *C. thermophila*, *E. monterogeii* and *Z. costaricensis*. Several data analysis features of the Phyloquant approach were tested: i) MS/MS identification and quantification software (Proteome Discoverer (PD) and MaxQuant (MQ)), ii) cut-off criteria for valid values identified (two and three valid values in at least one species, and 70% valid values in total; iii) protein and peptide normalized intensities from PD and LFQ from MQ; iv) normalized and non-normalized iBAQ intensities, and v) the application (or not) of the match-between runs (MBR) node in MQ (**Figure 1**). These criteria were first evaluated by principal component analysis, which allowed us to examine the clustering of the total proteomes prior to statistical analysis and subsequent evolutionary inferences.

**Figure 1.**
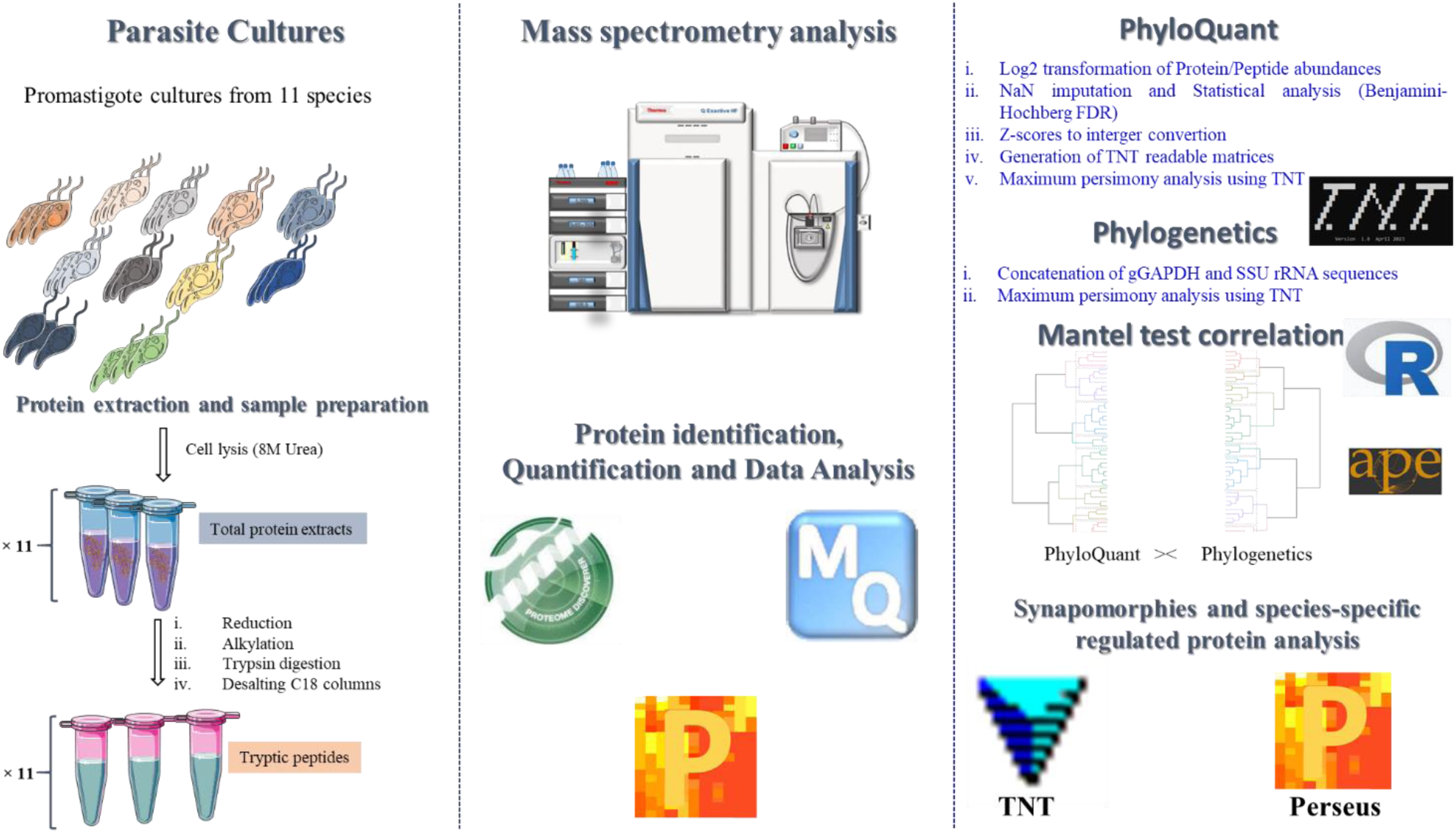
PhyloQuant analytical workflow applied to trypanosome species of the subfamily Leishmaniinae. Exponential phase promastigotes were cultured in TC100 medium, and the proteins extracted in 8M Urea, reduced in DTT, and alkylated using IAA before overnight trypsin digestion. The resulting tryptic peptides were desalted and submitted to nLC-MS/MS analysis, followed by protein identification and quantification using Proteome Discoverer (PD) and MaxQuant (MQ) softwares. Normalized protein and peptide intensities (PD), normalized iBAQ and LFQ protein intensities (MQ) were used to perform PhyloQuant clustering. Phylogenetic analysis using combined gGAPDH and SSU rRNA sequences was performed to evaluate the correlation with PhyloQuant using Mantel test. Protein expression profiles supporting each monophyletic cluster were mapped with TNT and visualized by Euclidean distance clustering using Perseus.

The number of identified and quantified proteins and/or peptides from PD (**Supplementary Table 1a-g**) and MQ, with (**Supplementary Table 2a-d**) and without (**Supplementary Table 3a-d**) MBR, and using different cut-off values for valid values of the identified proteins are summarized in **Supplementary Figure 1**. Proteome Discover software identified and quantified 16.7% more proteins and 19.9% more peptides (11634 and 69676, respectively) compared to MQ software using MBR (9693 and 55782, respectively). Less Non-assigned Numbers (NaNs) values were identified in PD compared to MQ analysis, as demonstrated after filtering for 70% valid values in total (**Supplementary Figure 1**), with 7048 and 211 proteins being identified after applying 70% of valid values filter in PD and MQ, respectively. Matrices of protein intensities from PD and MQ (iBAQ and LFQ) were used to infer evolutionary relationships between the different species under study. For PD, the analysis of the matrix from 70% valid values was possible due to the fewer NaNs compared to MQ. The difference between applying MBR or not in MQ searches was negligible, with the identification and quantification of 76 and 38 more proteins by iBAQ and LFQ intensities with MBR, respectively.

### Species-specific proteome expression patterns of species in the subfamily *Leishmaniinae*

Total proteome and peptide variability among the analysed species of the subfamily *Leishmaniinae* were evaluated using Principal Component Analysis (PCA). Log2 transformed intensities of normalized abundances of proteins and peptides from PD (**Supplementary Table 1**), proteins, and peptide LFQ intensities from MQ (**Supplementary Table 2,3**), and protein iBAQ intensities from MQ (**Supplementary Table 4**) with and without MBR were visualized by the first and second principal components of a PCA plot (**Figure 2**). For this analysis, imputation of NaNs was performed from normal distribution using a down-shift of 1.8 and distribution width of 0.3 before plotting the PCA plots.

**Figure 2.**
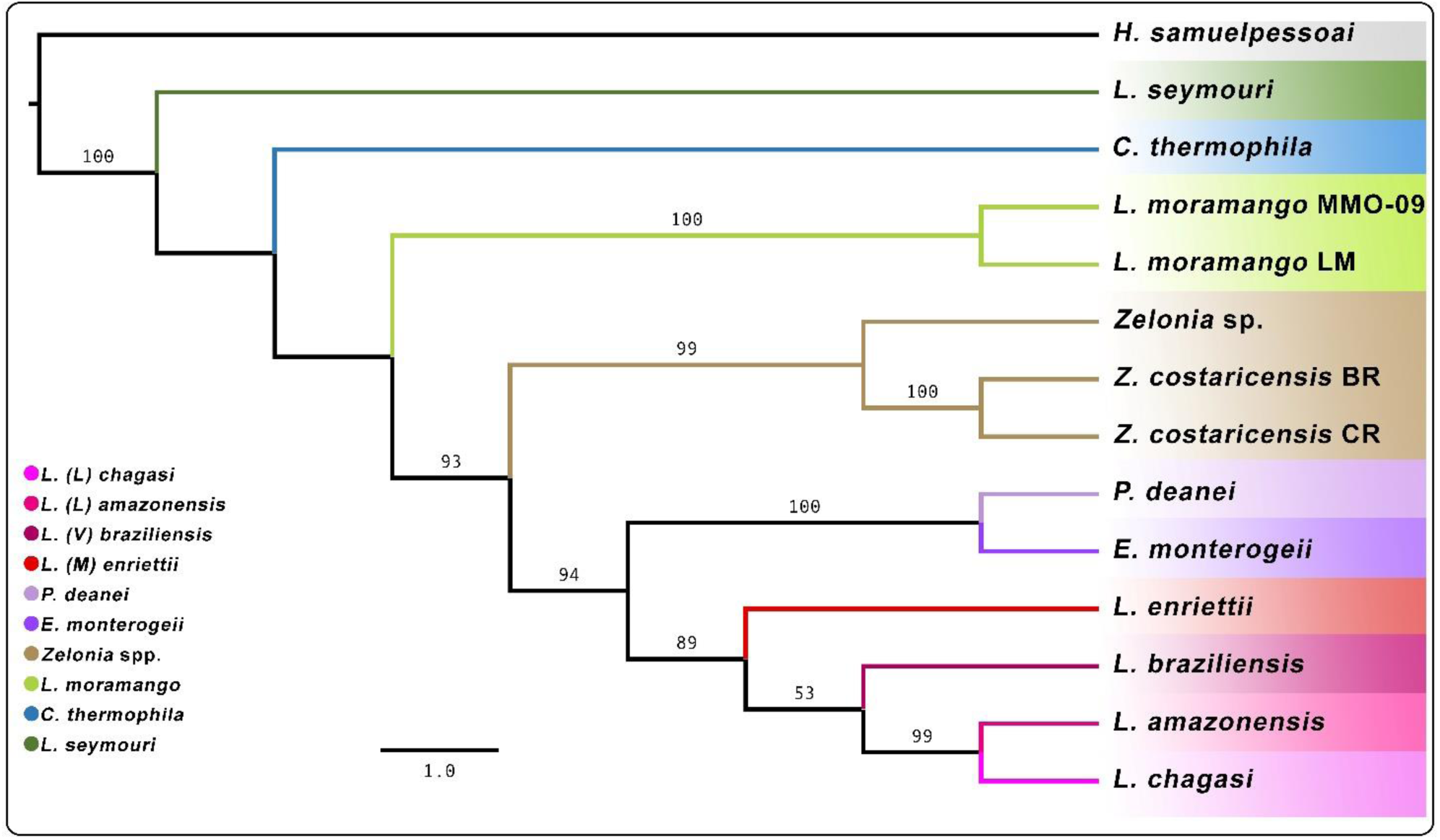
Phylogenetic relationships among trypanosomatids of the Leishmaniinae. gGAPDH and SSU rRNA concatenated DNA sequences from 11 species of six genera were used to infer the phylogenetic relationships by maximum parsimony (MP). Numbers at the major nodes represent MP support values (> 50) from 1000 replicates.

**Supplementary Figure 1.**
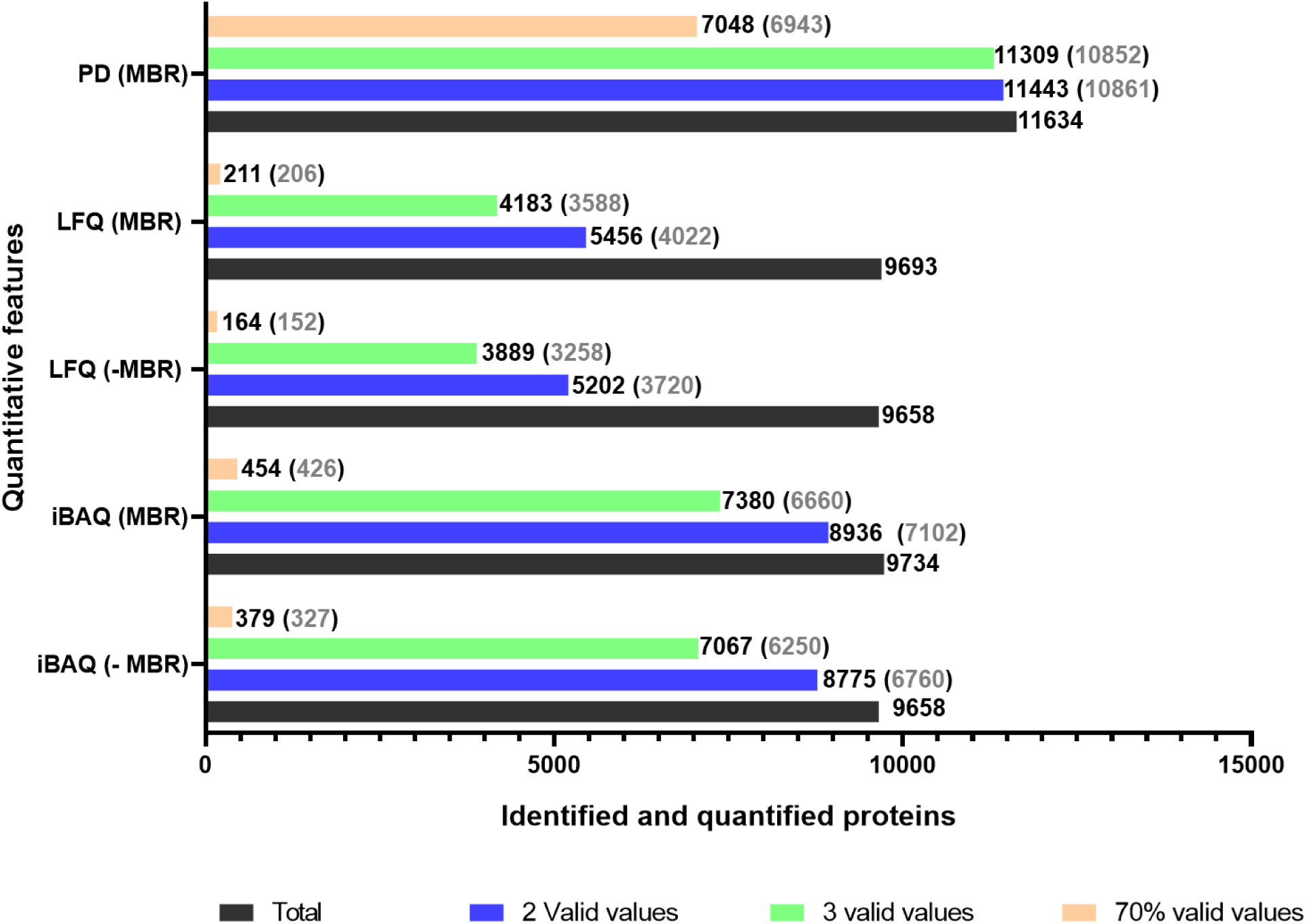
Total identified and quantified proteins from MaxQuant (MQ) and Proteome Discoverer (PD) softwares. The total number of proteins (grey), and proteins identified and quantified after applying a cut-off of two (blue) and three (green) valid values in at least one species, and 70% valid values in total (orange) are shown for MQ and PD, with the number of regulated proteins denoted in parenthesis. Both MQ searches with match between runs (MBR) and without match between runs (-MBR) for iBAQ and LFQ-based identification and quantification are shown.

Overall, the species belonging to *L. (Leishmania)* and *L. (Viannia)* subgenera clustered together and separately from the other trypanosomatids. Based on total proteome and peptide intensities from PD, *L. (M.) enriettii* was placed further from the assemblage comprising the other species of *Leishmania* (**Supplementary Figure 2 a, b**). However, *L. (M.) enriettii* was placed closer to the subgenera *L. (Leishmania)* and *L. (Viannia)* based on proteome and peptide intensities from LFQ and iBAQ, being closer to *L. (Viannia)* than *L. (Leishmania)* (**Supplementary Figure 2c, d, e, f**). The two species of *Zelonia*, *Z. costaricensis* (TCC 169E) and *Zelonia* spp. (TCC2547) clustered tightly together in all PCA analysis using PD, LFQ and iBAQ intensities. In addition, *P. deanei* and *E. monterogeii* also clustered tightly together in all analyses. *L. moramango* always clustered with *Zelonia* spp., and more related to *C. thermophyla* than to *L. seymouri* that was placed furthest from all species based on either the first (**Supplementary Figure 2c, d**) or second (**Supplementary Figure 2 a, b, e, f**) components.

**Supplementary Figure 2.**
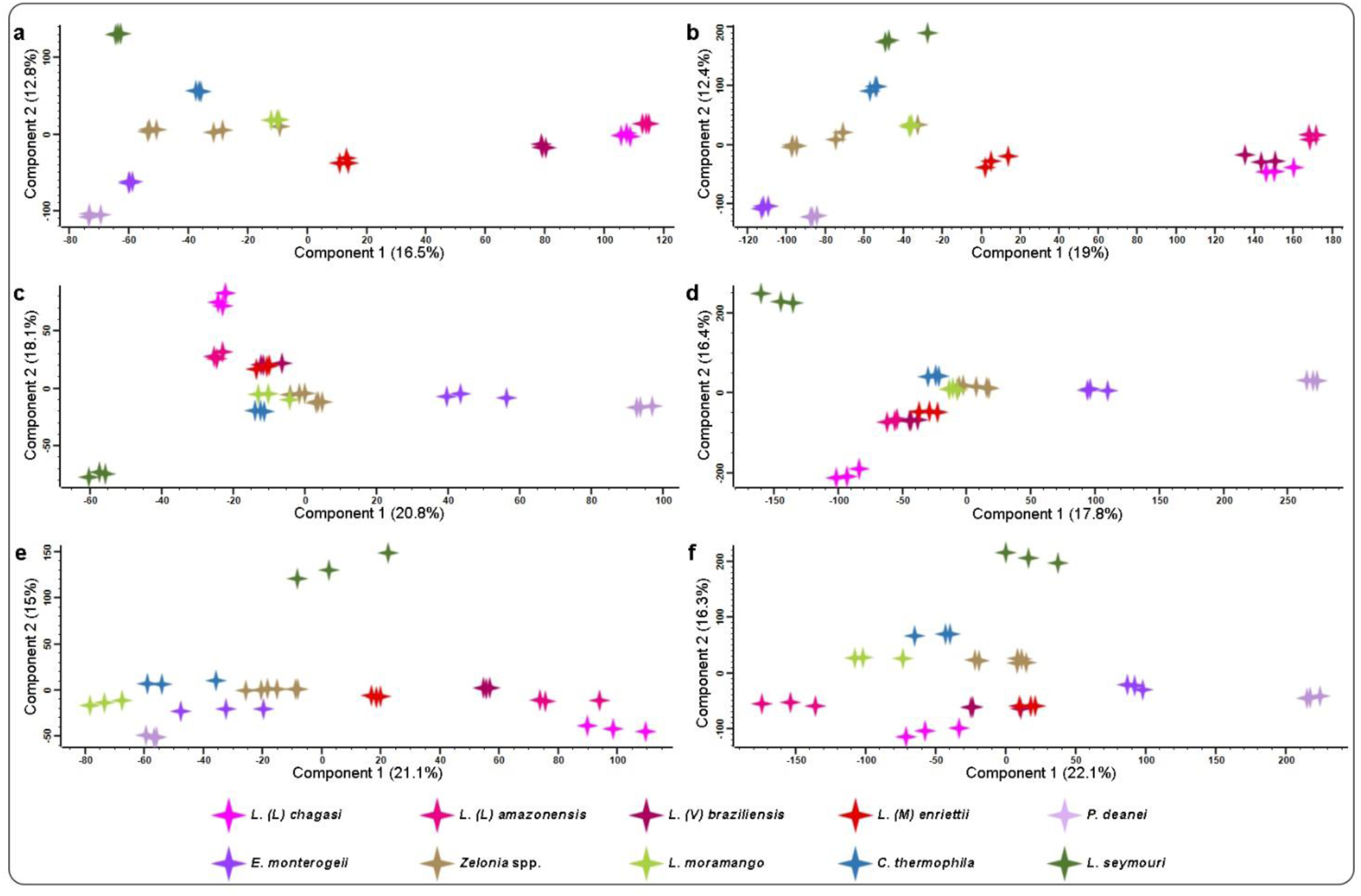
Principal component analysis of proteome and peptide variability among the analysed species of the subfamily Leishmaniinae. PCA clustering of trypanosomatids in the subfamily *Leishmaniinae* based on A) proteome and B) peptide variability using normalized abundances from Proteome Discoverer (PD) analysis; C) proteome and D) peptide LFQ intensities variability from MaxQuant (MQ) analysis; and proteome variability based on E) normalized and F) non-normalized iBAQ intensities from MQ analysis. In all analyses, three valid values in at least one species were applied as a cut-off, and the non-annotated numerals (NaNs) were imputed from a normal distribution.

### Phylogenetic resolution of several representative trypanosome species in the Leishmaniinae subfamily

Phylogenetic analysis was performed based on SSU rRNA and gGAPDH DNA sequences obtained from GenBank and TCC-USP data bank (**Figure 2**) on several species in the Leishmaniinae subfamily. This analysis showed the taxonomic resolution within the subfamily *Leishmaniinae* [4, 5], clearly separating *L. seymouri, C. thermophila* and *L. moramango* from the clade comprised of *Z. costaricensis* and *Zelonia* sp. TCC 2547, and from the major clade formed by [*Endotrypanum-Porcisia*] and [*Leishmania* spp.]. Also consistent with previous studies, *L.* (*M*.) *enriettii* was placed basal within the clade nesting *Leishmania* spp. of subgenera *Leishmania* and *Viannia*. The positioning of *L. moramango* separated from *L. seymouri* was concordant with phylogenetic inferences including a comprehensive sampling of species from both *Crithidia* and *Leptomonas* [31].

### PhyloQuant-based evolutionary inferences based on different quantification softwares and quantitative features

The influence of peptide and protein abundances were assessed by different identification and quantification softwares; PD and MQ, and used to evaluate the performance of the PhyloQuant method in resolving closely related taxonomic clusters. Taxonomic resolution and evolutionary inference of several species in the *Leishmaniinae* subfamily based on normalized protein and peptide abundances from PD software (**Figure 3**) formed five major clades in similar tree topologies. *L. seymouri* was always placed as the most basal species, supported by high support values (100), followed by *C. thermophila* and *L. moramango*; these three species never formed a clade. *L moramango* was placed at the edge of the major clade comprising *Zelonia*, *Endotrypanum* and *Porcisia* (**Figures 3a,b,c**). The two species of each genera clustered together, except *Zelonia* sp. TCC2547 which was placed closer to Endotrypanum/Porcisia in the analysis based on peptide intensities analysis (**Figure 3c**). The relationships between the sister clades formed by [*Endotrypanum*, *Porcisia* and *Zelonia*] and [Leishmania] is not well resolved. In contrast, in all three phylogenetic inferences based on differentially expressed protein abundances, all species of *Leishmania* clustered closely together. *L. (M.) enriettii* was basal to the other Leishmania species, more related to *L. (V.) braziliensis* than to the other species of the subgenera *Leishmania*, and those relationships were supported by high support values (99-100).

**Figure 3.**
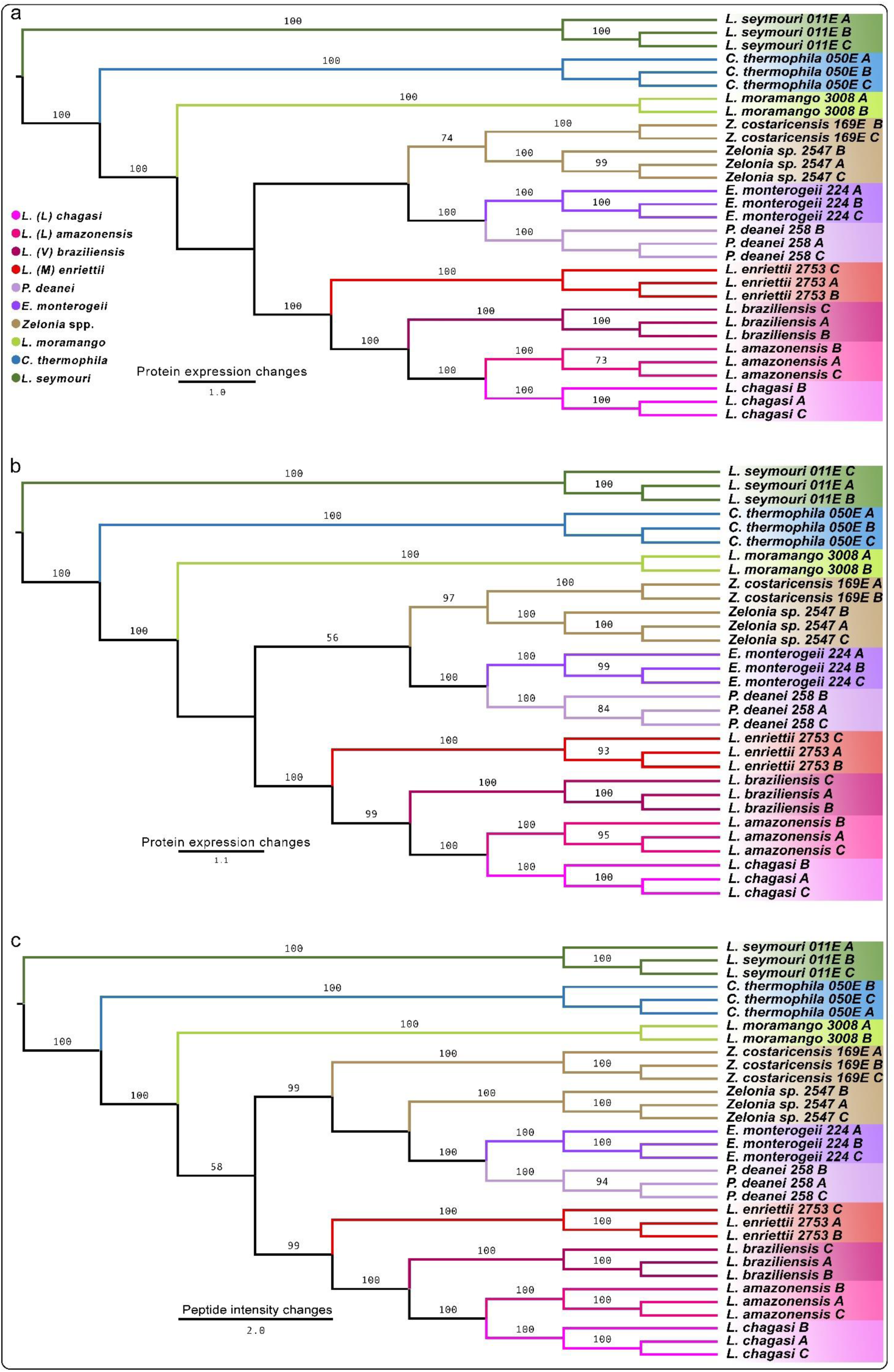
Evolutionary inference of representative species in the subfamily Leishmaniinae based on regulated protein and peptide abundances from Proteome Discoverer search engine. PhyloQuant approach on regulated normalized protein levels from a) three valid values in at least one species, b) 70% valid values in total, and c) peptide abundances from PD search engine. Phylogenetic analysis was performed using maximum parsimony in TNT for the three replicates for each species except for *L. moramango*. Branch support values from 1000 replicates are indicated.

LFQ values of protein and peptide expression intensities from MQ were also evaluated for the species from six genera nested into the *Leishmaniinae* subfamily using the PhyloQuant approach. Three and two valid values cut-offs were applied to the total identified and quantified proteins and peptides, respectively. Using differentially expressed proteins (3588) and peptides (25842) based on LFQ intensities, we show the high taxonomic resolution of the species, and their evolutionary inferences (**Figure 4**). Consistent with the previous analysis (**Figure 3**), *L. seymouri* was the most basal species, followed by *C. thermophila* and *L. moramango.* These species were always separated from the genera *Endotrypanum*, *Porcisia* and *Leishmania*, whereas *Zelonia* spp. were more closely related to *Porcisia/ Endotrypanum* than to the other monoxenous species. A major clade comprised of the sister groups formed by species of the *Zelonia* and the clade *P. deanei/E. monterogeii* were supported by protein and peptide LFQ intensities, supported by low (32%) and high (100%) bootstrap values, respectively. A second major clade comprising all species of the *Leishmania* subgenera was also supported by protein (79%) and peptide (100%) LFQ intensities, clustering *L. (M.) enriettii* and *L. (V.) braziliensis* as a group sister to the clade formed by *L. (L.) amazonensis* and *L. (L.) chagasi*. The relationship between the two major clades was supported by high (96%) and low (15%) bootstrap values for protein (**Figure 4a**) and peptide (**Figure 4b**) LFQ intensities, respectively.

**Figure 4.**
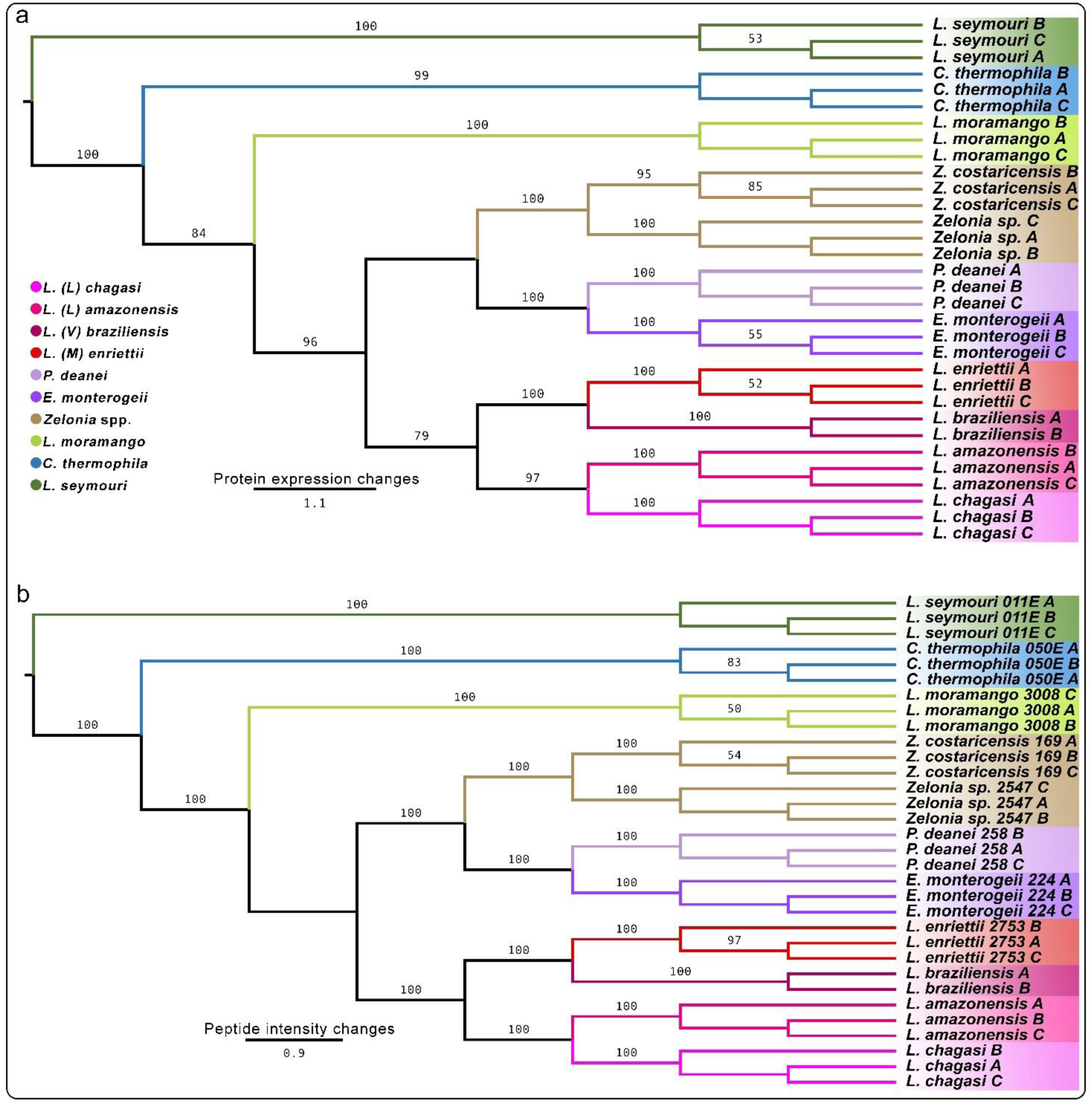
Evolutionary inference of Leishmaniinae species based on regulated protein and peptide LFQ intensities from MaxQuant search engine using the match between runs algorithm. Inference based on regulated a) protein and b) peptide LFQ intensities from MQ. Phylogenetic analysis was performed using maximum parsimony in TNT with three replicates of each species except for *L. (V.) braziliensis* replicate c. The branch support values from 1000 replicates are indicated.

Intensity-based absolute quantification (iBAQ) intensities from MQ searches with the MBR (**Supplementary Table 4a,b**) option were also evaluated for evolutionary inference using the PhyloQuant approach. In this analysis, differentially expressed proteins based on iBAQ intensities were evaluated without (**Figure 5a**) and with normalization (**Figure 5b**). Overall, the topologies of inferred relationships were consistent with those inferred in previous analyses (**Figures 3**, **4**). With *L. seymouri* as the most basal species, both quantitative MS features showed the separation of *C. thermophila* and *L. moramango* from *Zelonia*, *Endotrypanum*, *Porcisia,* and *Leishmania.* A high bootstrap value supported this separation based on non-normalized intensities (98%) (**Figure 5a**), while only a 2% support value was seen from normalized iBAQ intensities (**Figure 5b**). The clade comprised of *Zelonia* spp., *P. deanei* and *E. monterogeii* was maintained (96% support value) in the analysis based on non-normalized iBAQ intensities (**Figure 5a**), as previously shown using PD and LFQ abundances (**Figure 3** and **4**, respectively). However, based on normalized iBAQ intensities, the species of *Zelonia* formed a clade sister to that comprising *P. deanei* and *E. monterogeii*, supported by very low 2% bootstrap value (**Figure 5b**). As with LFQ intensities, the clade comprised of *L.* (*M*.) *enriettii* and *L. (V.) braziliensis* (supported by 100%) was separated from *L. (Leishmania)* based on non-normalized and normalized iBAQ intensities.

**Figure 5.**
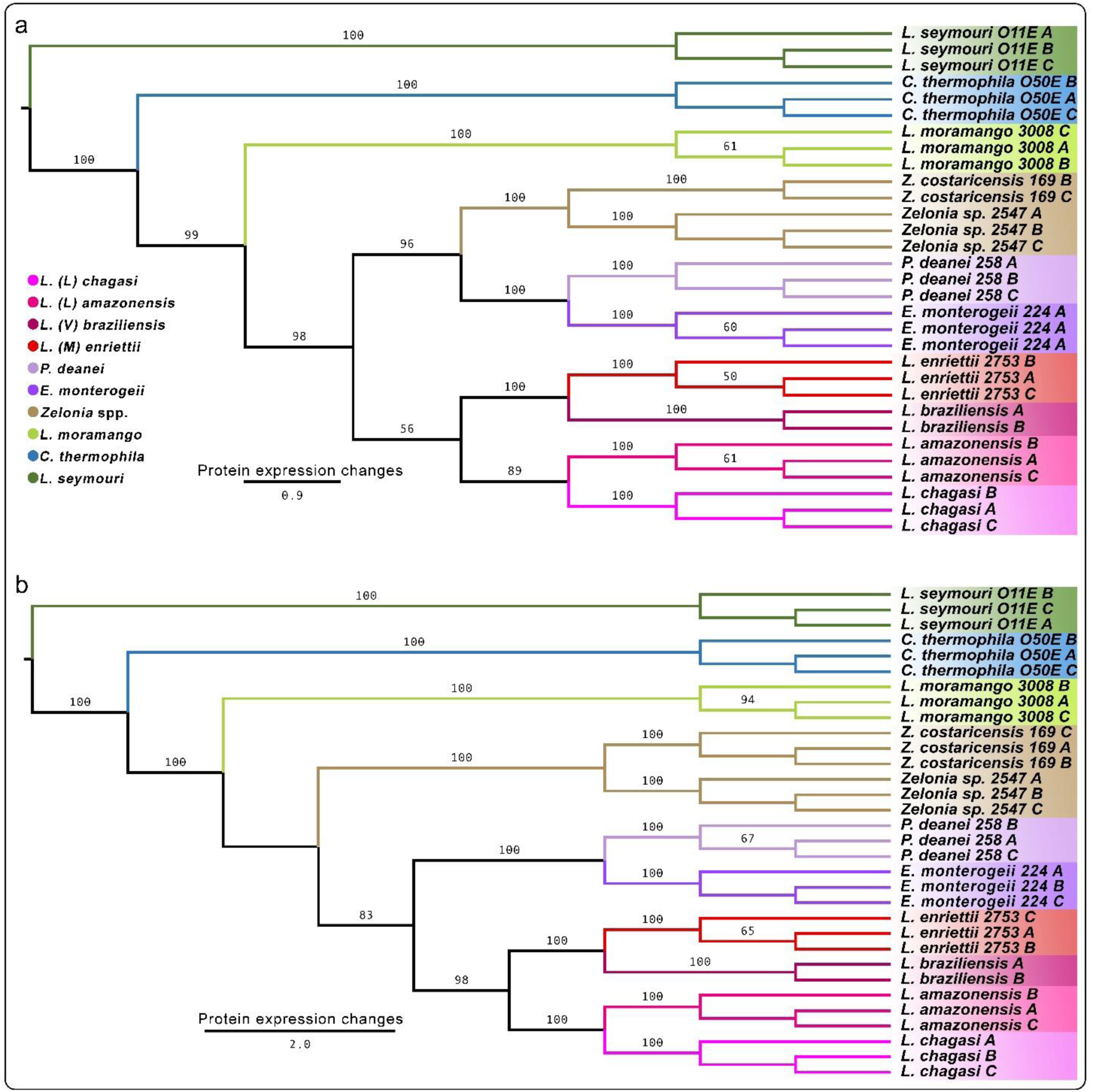
Evolutionary inference of representative species in the subfamily Leishmaniinae based on statistically significant proteins based in iBAQ intensities from MaxQuant search engine. PhyloQuant clustering based on statistically significant a) non-normalized iBAQ and b) normalized iBAQ intensities. Phylogeny analysis was performed using maximum parsimony in TNT for the three replicates for each species. Branch support values from 1000 replicates are indicated.

Moreover, MaxQuant LFQ intensities were searched without activating the match-between-runs node for comparisons with previous analysis (**Fig. 4**). LFQ intensities from identified and quantified proteins and peptides were used for evolutionary inferences (**Figure 6**). In both analyses, *L. seymouri*, *C. thermophila* and *L. moramango* were positioned separately in the same order inferred by previous analyses (**Figures 3,4,5**), and separated from *Zelonia*, *Endotrypanum*, *Porcisia* and *Leishmania* (**Figure 6a, 6b**). The major clade comprised of *Zelonia* and *P. deanei/E. monterogeii* was supported by 100% and 7%, while the other clade nesting all *Leishmania* spp. were always strongly supported (88-98%). As before, discordant values ranging from 53% to 1% supported the relationship between these two major clades based on proteins and peptide abundances (**Figures 6a** and **6b**, respectively). The relationship among the *Leishmania* subgenera was supported by high branch support values based on both protein (88%) and peptide (98%) LFQ intensities (**Figures 6a, 6b**).

**Figure 6.**
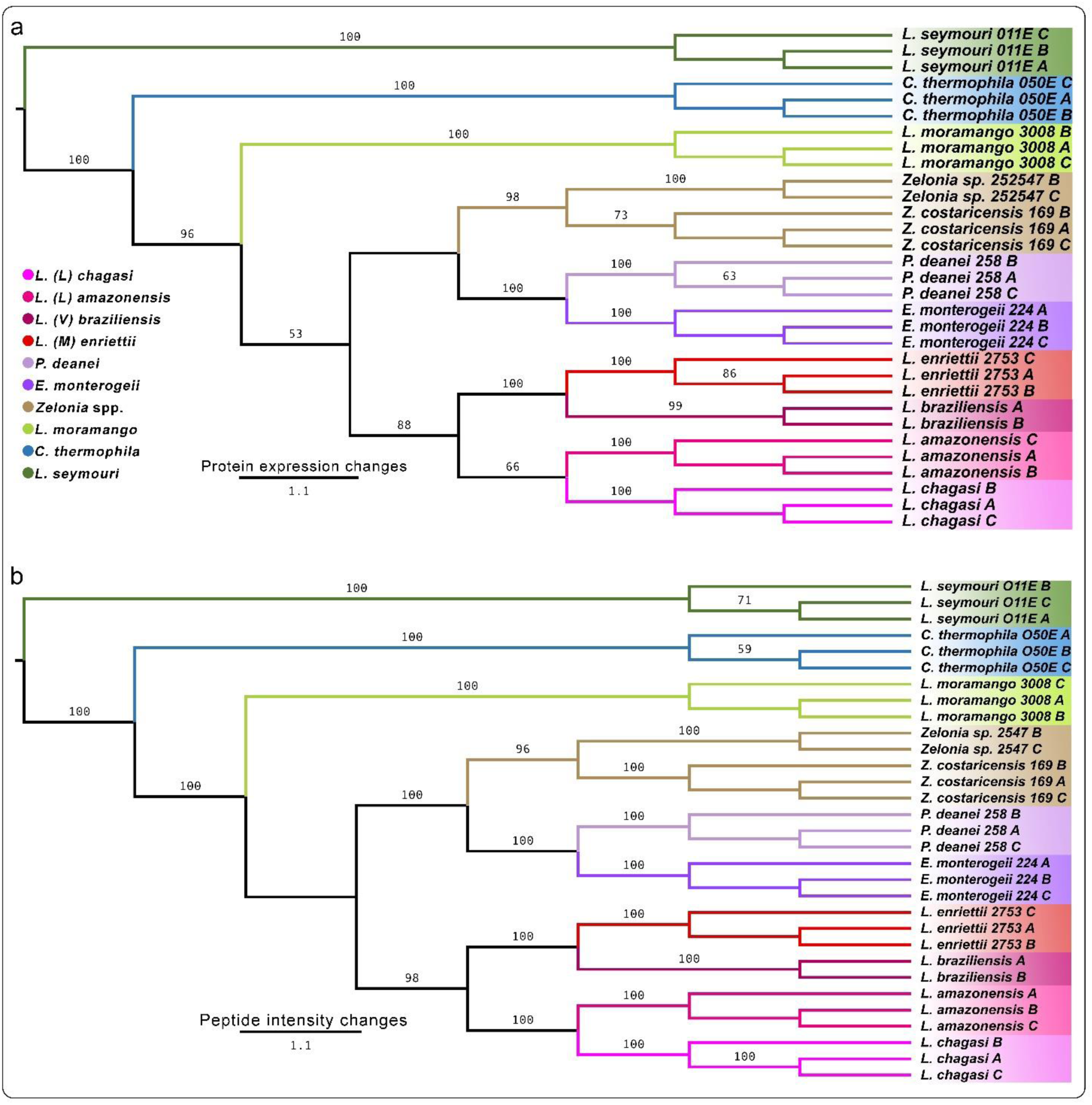
Evolutionary inference of representative species in the subfamily Leishmaniinae based on statistically significant LFQ intensities of proteins and peptides from MaxQuant search engine without the Match between runs algorithm. PhyloQuant clustering based on statistically significant a) protein and b) peptide LFQ intensities. Phylogeny analysis was performed using maximum parsimony in TNT for the three replicates for each species. Branch support values from 1000 replicates are indicated.

### Evolutionary congruence between Leishmaniinae Phylogenetics and PhyloQuant

Mantel test was performed to calculate the levels of correlation between sequence-based phylogenetic tree (**Supplementary Figure 3**) and PhyloQuant-based phylogenies based on the different mass spectrometry-based quantitative features and quantification platforms. One representative replicate for each trypanosome was used to infer evolutionary relationships based on PhyloQuant approach before performing Mantel test for correlation. The *p*-values and Z-statistic values from the Mantel test are summarized in **Table 2**. All evolutionary trees based on the PhyloQuant method showed a statistically significant level of correlation (*p*-0.0009) with the sequence-based phylogenetic tree, with high Z-statistic values. Random trees with similar nodes were included in this analysis, showing that only two of the ten random trees (tree 1 and tree 6) were significantly correlated to the phylogenetic tree generated by MP using concatenated V7V8 SSU rRNA and gGAPDH gene sequences. Correlation of the phylogenetic tree matrix to itself yielded the highest Z-statistic values, and the same statistical significance as all the evolutionary trees based on PhyloQuant analysis. Trees based on normalized peptide and protein intensities from Proteome Discover, and normalized iBAQ from MBR searches gave high Z-statistic values.

**Table 2.** Mantel test correlation between the PhyloQuant approach, based on different mass spectrometry-based intensities, and phylogenetic analysis based on SSSU rRNA and gGAPDH gene sequences. PD = Proteome Discoverer; MBR – Match Between Runs; −MBR – No Match Between Runs. The evolutionary trees and the supplementary tables with Log2 protein/peptide expression abundances used to generate the trees are also illustrated.

| Evolutionary tree | Figure | Supplementary Table | Z statistic | p-value |
| --- | --- | --- | --- | --- |
| Phylogenetics | Sup. Fig. 3 | - | 7340 | 0.000999 |
| PD_Proteome | Fig. 3a | 1c | 6176 | 0.000999 |
| PD_Proteome, 70% valid values | Fig. 3b | 1e | 6176 | 0.000999 |
| PD_Peptide | Fig. 3c | 1g | 6224 | 0.000999 |
| LFQ_Proteome, MBR | Fig. 4a | 2b | 5076 | 0.000999 |
| LFQ_Peptide, MBR | Fig. 4b | 2d | 5076 | 0.000999 |
| iBAQ_non-normalized, MBR | Fig. 5a | 4a | 5076 | 0.000999 |
| iBAQ_normalized, MBR | Fig. 5b | 4b | 6160 | 0.000999 |
| LFQ_Protroteome, -MBR | Fig. 6a | 3b | 5076 | 0.000999 |
| LFQ_Peptide, -MBR | Fig. 6b | 3d | 5076 | 0.000999 |
| random tree1 | - | - | 2029 | 0.000999 |
| random tree2 | - | - | 1666 | 0.068931 |
| random tree3 | - | - | 1687 | 0.098901 |
| random tree4 | - | - | 1058 | 0.374625 |
| random tree5 | - | - | 1692 | 0.042957 |
| random tree6 | - | - | 1392 | 0.004995 |
| random tree7 | - | - | 1507 | 0.152847 |
| random tree8 | - | - | 1548 | 0.576424 |
| random tree9 | - | - | 1824 | 0.182817 |
| random tree10 | - | - | 1386 | 0.206793 |

Evolutionary inference based on PD data showed the placement of *L. (Mundinia*) subgenus, here represented by *L. (M.) enriettii,* as the basal clade to the other *Leishmania* subgenera, which was corroborated by the DNA sequence-based phylogenetic tree. Evolutionary inference based on normalized iBAQ intensities (**Figure 5b**) showed the separation of *Zelonia* from the *Porcisia*-*Endotrypanum* clade, also observed in the phylogenetic analysis. These tree topologies would support the higher Z-statistic values compared to other tree topologies which clustered *L. braziliensis* sister to *L. (M.) enriettii*, and the clade *Porcisia*-*Endotrypanum* sister to *Zelonia* differing from phylogenetic tree topology.

**Supplementary Figure 3.**
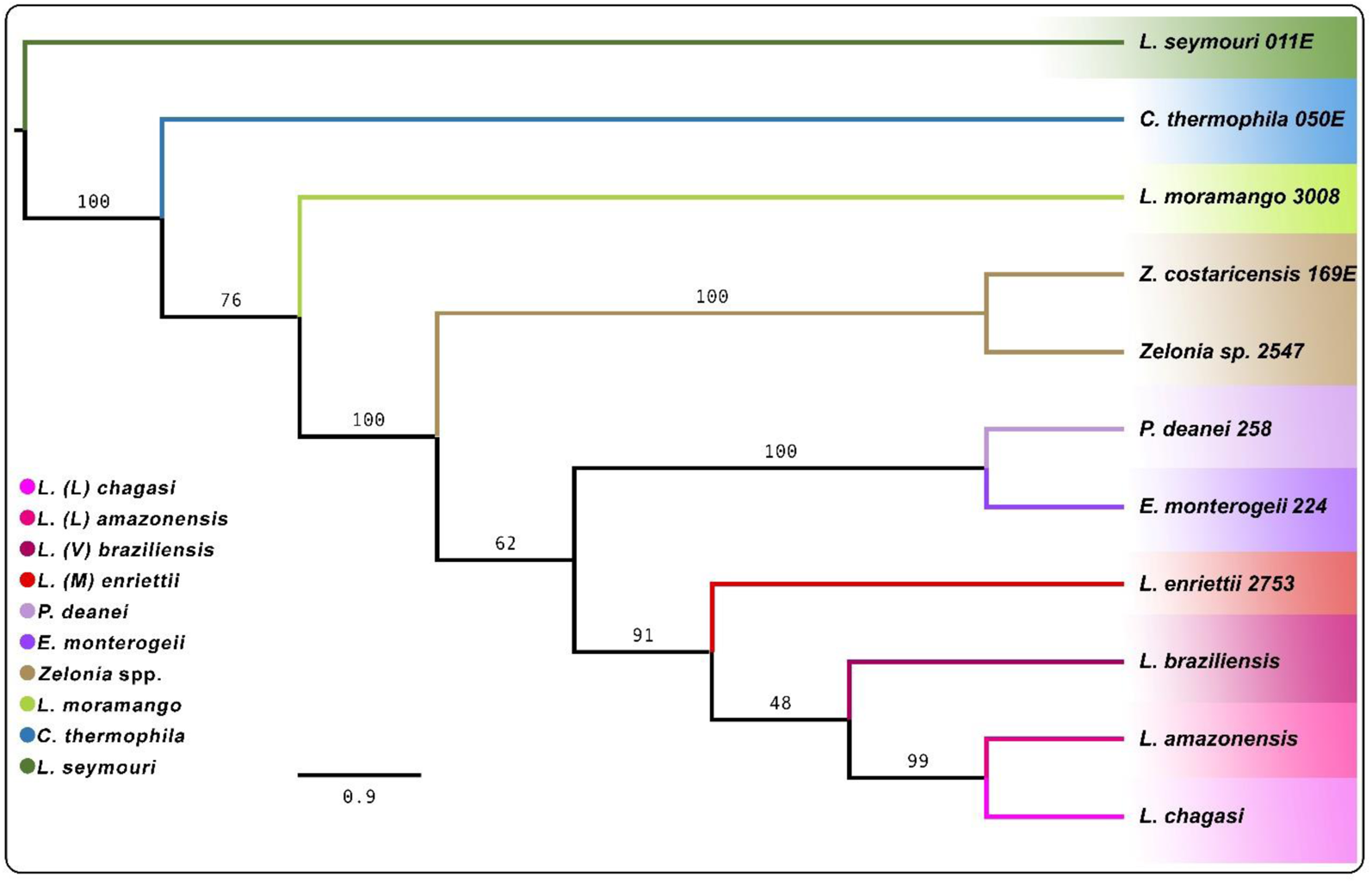
Phylogenetic tree of several trypanosomatid species in the Leishmaniinae subfamily previously evaluated in the PhyloQuant analysis for mantel test correlation. gGAPDH and SSU rRNA concatenated sequences were used to infer the phylogenetic relationships using Maximum Parsimony. Numbers at the major nodes represent MP support values from 1000 replicates.

**Supplementary Figure 4.**
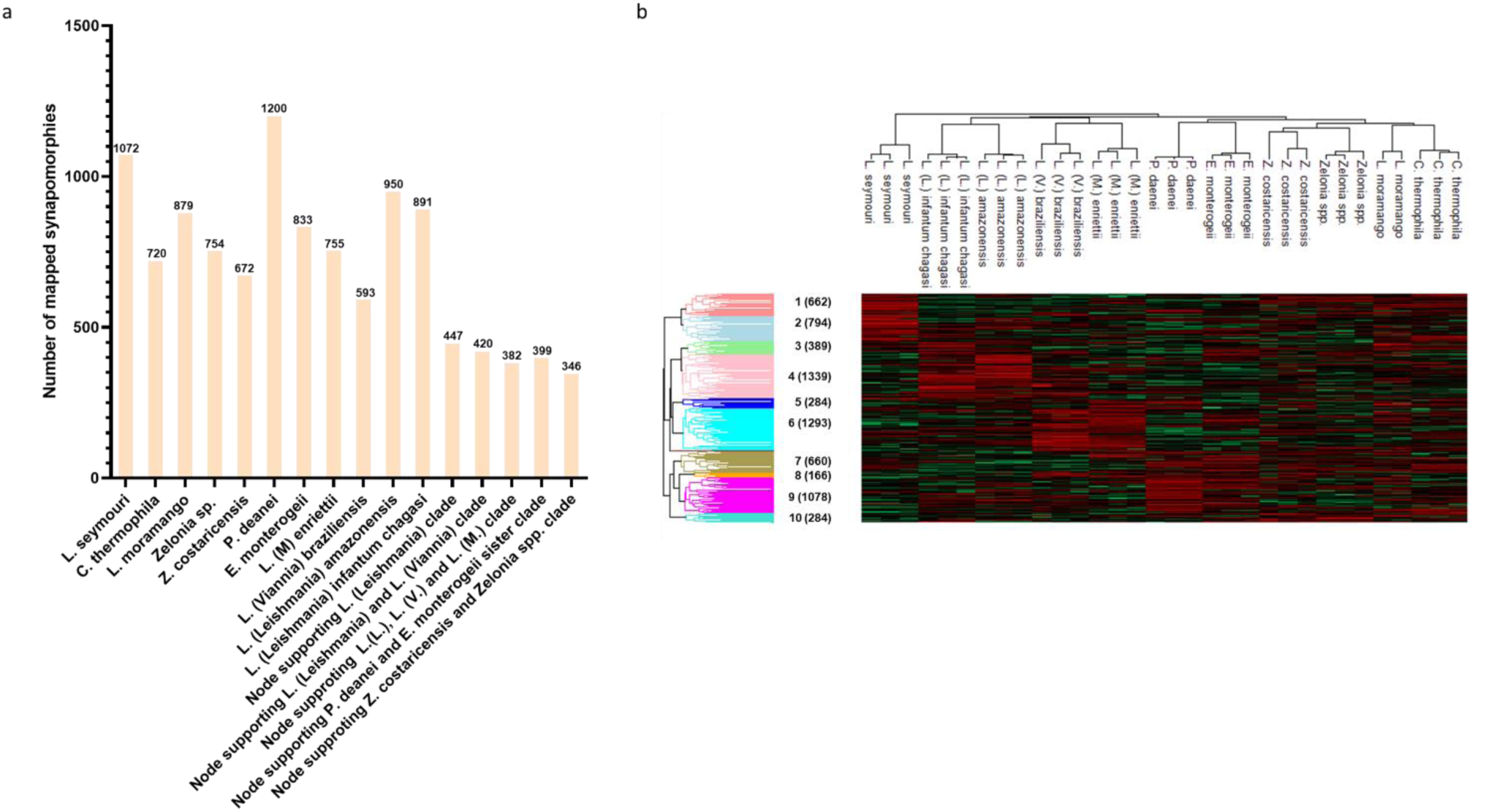
Number of mapped synapomorphies and Euclidean distance-based clustering of species in the subfamily Leishmaniinae based on differentially expressed proteins using PD software. a) Number of mapped synapomorphies for the different species and clusters, b) Clusters of upregulated and downregulated proteins for each species are illustrated in red and green, respectively.

### Synapomorphies and species-specific upregulated protein expression profile analysis

Based on the regulated proteins identified and quantified by the Proteome Discoverer with a cut-off of 70% valid values, we mapped the synapomorphies supporting the major clades using TNT software (**Supplementary Table 5**), and the upregulated proteins for the different species using Euclidean clustering (**Supplementary Table 6**). The synapomorphies and their expression profiles are summarized in **Supplementary Figure 4a**, and the heatmap following Euclidean distance clustering showing the species-specific protein expression profiles in **Supplementary Figure 4b**. *L. seymouri* and *P. deanei* had the highest numbers of mapped synapomorphies (1072 and 1200), while the expression profiles of 879 and 720 proteins supported the polyphyletic clusters of *C. thermophila* and *L. moramango*, respectively. Interestingly, fewer number of proteins supported *L. braziliensis* (593) clade, which was separated from *L. amazonensis* and *L.* (*L.*) *infantum chagasi*, supported by 950 and 891 proteins, respectively. A total of 382 synapomorphies support the *Leishmania* genus (*L*. (*Leishmania*), *L*. (*Viannia*) and *L. mundinia*), while the sister clades [*P. deanei* and *E. monterogeii*) and [*Zelonia* sp. and *Z. costaricensis*] are supported by 399 and 346 synapomorphies, respectively.

In addition to clustering the species according to resolved taxons, we used Euclidean distance clustering to visualize species-specific regulated protein profiles (**Supplementary Figure 4b**; **Supplementary Table 6**). Cluster 1, with 662 proteins, comprised of proteins which were upregulated in *L. seymouri*, Zelonia spp. and *C. thermophila*, while cluster 2, with 794 proteins, represents upregulated proteins in *L. seymouri* and *L. moramango*. Proteins specifically upregulated in *L.* (*L.*) *infantum chagasi* are represented by cluster 3 (389 proteins), and cluster 4, with 1339 proteins, represents upregulated proteins in *L.* (*L.*) *infantum chagasi*, *L.* (*L.*) *amazonensis* and *L.*(*V.*) *braziliensis*. Clusters 5 and 6, with 284 and 1293 proteins, respectively, highlight proteins upregulated in *L. braziliensis* and *L. (M.) enriettii*, while cluster 7 (660 proteins) represents proteins which higher abundances in *P. deanei*, *E. monterogeii* and *C. thermophila*. Cluster 8 shows 166 upregulated proteins in *L. (V.) braziliensis*, *L. (M.) enriettii* and *P. deanei*, while cluster 9 shows proteins specifically more abundant in *P. deanei*. Cluster 10 has 284 proteins with higher expression intensities shared by both *Zelonia costaricensis* and *Zelonia* sp.

## Discussion

In the Trypanosomatidae family, monoxenous life cycle is ancestral to dixenous life, and the comparison between trypanosomatids from insects and those that developed in both vertebrate and insect (vector) hosts are crucial for the understanding on their origin, host-switching from insects to vertebrates, adaptation to new hosts and vectors, and the emergence of pathogens. In the present study we conducted a PhyloQuant approach to assess differential protein expression profiles to gain insights into the phylogenetic relationship among the species of the *Leishmaniinae* subfamily, by comparing protein profiles of 11 species from six genera. This subfamily led by *Leishmania* was created to accommodate trypanosomatids of *Endotrypanum, Leptomonas, Crithidia,* and *Novymonas* [3]. Recent studies have revealed new species, subgenera, and genera within Leishmaniinae: the subgenus *Mundinia*, the new genera *Zelonia* and *Porcisia* [4, 5, 32], and the genus *Borovskyia,* with a single species found in Costa Rica [5, 33].

For the present study, we selected six dixenous species, including reference-species of the three subgenera of *Leishmania* comprising human pathogens (*Leishmania*, *Viannia* and *Mundinia*), *Endotrypanum* and *Porcisia.* These species were compared with five monoxenous species of *Zelonia*, *Leptomonas* and *Crithidia*. To date, only species of *Leishmania* and *Endotrypanum* were reported causing human infections, all transmitted by sand flies. The genus *Endotrypanum* is characterized by intraerythrocytic trypomastigotes and/or epimastigotes in sloths. However, *E. colombiensis* is known to cause both cutaneous and visceral leishmaniasis in humans. Intracellular amastigotes are obligatory in *Leishmania* and have been recorded for E. colombiensis in man. *Porcisia* in porcupines and *Endotrypanum* in sloths and squirrels apparently do not cause any disease symptoms [4, 13–17].

The present study included an isolate of *Z. costaricensis* from Brazilian Amazonia, which is identical to the first isolate of this species from Costa Rica, Central America [4] and a new species of *Zelonia* (TCC 2547) from Panama, Central America. Nested within Leishmaniinae, but more distant from *Leishmania* than *Endotrypanum, Porcisia* and *Zelonia*, were placed *Leptomonas* and *Crithidia*, comprising species from insects of Heteroptera, Diptera and Hymenoptera. The species classified in the genera *Crithidia* and *Leptomonas* cannot be reliably separated from each other, and both genera are polyphyletic as reinforced in the present study. Our study includes two distant related species classified as *Leptomonas*, but clearly belonging to different genera: *L. seymoury* and *L. moramango. L. moramango* isolates TCC3008 and MMO-09 [34] obtained from dipterans collected in Uganda and Madagascar, respectively, shared identical sequences and exhibited typical promastigotes. *L. seymouri* is predominantly monoxenous of hemipterans, while it can infect vertebrates, even humans, opportunistically. The genus *Crithidia,* type-species *C. fasciculata,* was herein represented by *C. thermophila,* reported from dipterans and hemipterans over the world. Both *L. seymouri* and *C. thermophila* displayed thermotolerance (thermal resistance up to 34°C), a prerequisite for living in warm-blooded vertebrates [5, 31, 35].

We compared promastigote forms from most species of all genera, except for *C. thermophila* that displays choanomastigote forms. Promastigote molecules mediate the differentiation of infective forms in vectors and are vital to parasite survival during early infection phase, playing vital roles in their phagocytosis and recognition in dendrocytes, neutrophils, and macrophages and immune evasion as well as contributing to virulence of *Leishmania* spp. [36–38].

Using several mass spectrometry-based quantitative features and different identification and quantification software, we show the consistency of the PhyloQuant approach to evolutionary inferences, thus providing additional support for taxonomy based on DNA-based phylogenetic relationships. Proteome Discoverer identified and quantified more proteins compared to MaxQuant, and with fewer NaN values. In addition, PhyloQuant analysis showed better resolution power and evolutionary relationships using normalized abundances from PD compared to MQ. PD-based PhyloQuant using both protein and peptide intensities placed *L. enriettii* (subgenera *L. (Mundinia*) in the edge of the clade nesting *L. (Leishmania)* and *L. (Viannia)* species. This clade was sister to that constituted by *Endotrypanum* and *Porcisia*, which together with *Leishmania* formed a monophyletic assemblage, whose parasitism in blood-sucking dipterans, such as phlebotomine sand flies and ceratopoginid midges, likely favour their adaptation to mammals. In most analyses, the two species of *Zelonia* clustered together and more closely related to *Porcisia/ Endotrypanum* than to any other monoxenous species. Although reported from blood-sucking black fly (Simulidae), *Zelonia* is so far restricted to insects [18]. *L. seymouri* was the most basal species, followed by *C. thermophila* and *L. moramango.* The topologies of phylogenetic inferences based on PhyloQuant were highly congruent with that showed by phylogenetic trees constructed using gene sequences.

Different proteomic approaches have been frequently employed to investigate *Leishmania* and leishmaniasis. High-throughput proteomics are commonly evaluated to identify protein targets for diagnosis and treatment, and to understand mechanisms involved in the infection and disease manifestations. In addition, proteome wide phylogenetic analyses have been explored to assess evolutionary processes in *Leishmania* spp. Proteome-wide phylogenomic analyses restricted to *Leishmania* species based on predicted protein sequences highlighted targets with potential involvement in virulence, pathogenicity, host-parasite interaction, and clinical manifestation of cutaneous and visceral leishmaniasis [39]. Previous studies focused on metaphylomes of *Leishmania* spp. revealed different protein repertoires related to subgenus, species and intra-specific diversification [39–41].

Evolutionary background that gave rise to specific adaptations responsible for the switch from monoxenous to dixenous life, and on the evolution of pathogenicity within Leishmaniinae is mostly limited to genome data, transcriptome, and predicted proteomes inferred from genomes. Proteome-wide comparative analyses between monoxenous and dixenous species of Leishmaniinae remains to be addressed. A few proteomic studies compared *Leishmania* and other Leishmaniinae genera. The comparison of human isolates of *L. infantum* and *L. seymoury* uncovered some proteins that can be related to worst disease and drug resistance induced by co-infections [42].

Most differential protein expression between *Leishmania* and monoxenous species were uncovered from transcriptome analyses. By comparing expression profiles of *L. mexicana*, *L. major* and *L. seymouri*, proteins potentially involved in virulence were identified. Additionally, different protein expression profiles were uncovered for *C. thermophila* and *L. seymouri,* which despite well adapted to warm-blooded hosts are unable of infecting macrophages and other mammalian cells, and apparently do not develop in sand flies [31, 35, 43]. A phylogram constructed by TOMM (Total Ortholog Median Matrix) approach using median amino acid distances between all protein-coding genes (∼5636 orthologous protein pairs across 46 kinetoplastida species) rather than selected proteins also supported the relationships among Leishmania, *Endotrypanum*, *Crithidia*, and *Leptomonas* [44]. Phylogeny based on 410 proteins encoded by single copy genes predicted from genomes corroborated *Endotrypanum* and *Porcisia* spp. as the closest known relatives of *Leishmania*, increased gene family losses and contractions compared to *Leishmania*, and reduced repertoire of surface proteins critical for *Leishmania* development within macrophages. Although it is considered that *Endotrypanum* and *Porcisia* do not appear to generally develop within macrophages *in vitro* [45], they are known to be intracellular parasites [15, 46, 47]. There is considerable genetic diversity within the poorly studied *Endotrypanum* clade [4] and it is feasible that these intracellular parasites employ alternative pathways to infect and multiply within their mammalian host’s cells.

## Conclusions

To provide new insights to phylogenetic relationships and reliable taxonomy of Leishmaniinae we carried out the most comprehensive proteome-wide analysis of this subfamily. The PhyloQuant approach supports current phylogeny and taxonomy of Leishmaniinae and corroborate a new species of Zelonia and the polyphyly of *Leptomonas*. As we previously demonstrated for trypanosomes of the subgenus *Schizotrypanum* [1], mass spectrometry-based quantitative features and the use of different identification and quantification software reinforced the consistency of the PhyloQuant approach to provide new information that can aid resolve evolutionary relationships for taxonomical purposes. In addition, this technique allows for identifying protein expression profiles differing according to subgenus/species, thus unveiling proteins that can be used to understand the differences in many biological processes, host-parasite-vector interactions, and infectivity, pathogenicity, virulence, and disease manifestations.

## Acknowledgements

The work was supported by grants and fellowships from FAPESP (2018/18257-1, 2018/15549-1, 2020/04923-0 to GP; 2017/04032-5 and 2021/14751-4 to SNM; 2016/23689-2 to JSS; 2018/13283-4 to GSDO, 2021/14179-9 to DMS, and 2020/13562-0 to MC), and from Conselho Nacional de Desenvolvimento Científico Tecnológico (CNPq) (“bolsa de produtividade” to GP). We would also like to acknowledge and thank Omar A. Espinosa for early studies on Zelonia spp. from Panama, and Marta Campaner for her support in culturing the parasites used in this study. Lastly, we would like to acknowledge the late Prof. Erney P. Camargo for the constructive and enlightening discussions during this study.

